# Comparison of image registration techniques in functional ultrasound imaging

**DOI:** 10.1101/2023.09.15.557999

**Authors:** Shan Zhong, Kofi Agyeman, Shanze Syed, Richard Tobing, Wooseong Choi, Charles Liu, Darrin Lee, Vassilios Christopoulos

## Abstract

Functional Ultrasound Imaging (fUSI) is an emerging hemodynamic-based functional neuroimaging technique that combines high spatiotemporal resolution and sensitivity, as well as extensive brain coverage, enabling a range of applications in both control and disease animal models. Based on power Doppler (pD) imaging, fUSI measures changes in cerebral blood volume (CBV) by detecting the back-scattered echoes from red blood cells moving within its field of view (FOV). However, the expansion of fUSI technology is partly limited by the challenge to co-register pD vascular maps acquired across different sessions or animals to one reference; an approach that could widen the scope of experimental paradigms and enable advanced data analysis tools. In this study, we seek to address this critical limitation. We evaluate six image registration techniques, predominantly used in other neuroimaging studies, using 2D sagittal whole-brain fUSI data from 82 anesthetized mice, and tested the quality of registration using multiple metrics. Our findings indicate a substantial enhancement in the alignment of fUSI images post registration. Among the tested techniques, the non-rigid registration algorithm *Imregdeform* yielded superior performance. We offer the first comparative study of image registration techniques for a 2D fUSI brain dataset, paving a way for improved utilization of fUSI in future pre-clinical research applications.

## 1 Introduction

Functional ultrasound imaging (fUSI) represents a rapidly advancing neuroimaging modality for large-scale recordings of neural activity through neurovascular coupling [1–4]. It is minimally invasive and provides a unique combination of great spatial coverage (*∼* 10 cm), high spatiotemporal resolution (*∼* 100 µm, up to 10 ms) and sensitivity (*∼* 1 mm/s velocity of blood flow). The enhanced spatiotemporal properties allow for sensing of the function of small neuronal populations, providing a closer connection to underlying neuronal signal compared to other hemodynamic methods, such as functional magnetic resonance imaging (fMRI). The first *in vivo* proof-of-concept for fUSI was established in 2011 by imaging the cerebral blood volume (CBV) changes in the micro-vascularization of the rat brain during whisker stimulation [2]. Since then, fUSI has been applied to image brain activity during olfactory stimuli [5], resting state functional connectivity [6], behavioral tasks in freely moving rodents [7], and non-human primates [8, 9], while recently it has also been extended to clinical studies [10, 11].

While fUSI has enabled a wide range of pre-clinical and clinical applications, the lack of fUSI-specific image registration techniques that enable effective alignment of signals into a reference power Doppler (pD) vascular template limits the scope of possible functional analysis. Bregma coordinates are utilized to establish precise and uniform location for fUSI 2D-plane imaging within, across acquisitions and across animals in pre-clinical brain imaging studies. However, the lack of effective alignment and normalization algorithms for fUSI images makes it challenging to perform random effect analysis and longitudinal experiments. Therefore, almost all fUSI studies are restricted to either averaging activity within selected regions of interest (ROIs) [9, 12, 13] or ROI-ROI functional connectivity analysis across sessions and animals [6, 14] (for an excellent review see [15]).

Given this context and the increasing adoption of fUSI for functional neuroimaging studies, there is a clear and distinct need to develop techniques for co-registering pD vascular maps acquired across multiple sessions and/or different animals. This will enable investigators to explore a wider spectrum of experimental paradigms, leverage more advanced machine learning tools, voxel-wise analysis and generate statistical parametric maps (SPMs) from data pooled from multiple sessions and across animals. In the current study, we evaluate the effectiveness of 6 distinct image registration techniques that have been widely used in neuroimaging studies. We employ 2-dimensional sagittal whole-brain fUSI pD intensity data acquired from anesthetized mice and demonstrate that the alignment of registered fUSI images are significantly improved over the alignment of pre-registration images (moving images). We extend our results to show that the non-rigid registration algorithm *Imregdeform* outperforms all other techniques tested. Overall, our study provides a comparison of image registration techniques for 2D fUSI brain data, promoting more effective use of fUSI in pre-clinical research settings.

## 2 Materials and Methods

### 2.1 Animals

The dataset used in the current study is part of another research project performed by our team recently to understand the mechanism of medial septal nucleus (MSN) deep brain stimulation (DBS) on CBV changes following MK-801 drug administration [16]. 82 male 8–12-week-old C57BL/6 mice (Charles River Laboratories; Hollister, CA) were used in this study. They were fed *ad libitum* and maintained at a regular light-dark cycle of 12h. Mice were anesthetized with 5% isoflurane in O_2_/N_2_O (1:2) carrier gas and then maintained at a constant rate (1.5-2%) through the experiment. Body temperature was kept constant throughout fUSI recordings by placing animals on an electric heating pad. The mice were then visually monitored for respiration and reflexes. Hair was removed from the animals’ head using a commercially available depilatory cream (Nair, Pharmapacks). They were then head-fixed in a stereotaxic frame with ear bars to stabilize their heads. Echographic gel was then applied onto the imaging window. All procedures were approved by the Institutional Animal Care and Use Committee of University of Southern California (IACUC #21006).

### 2.2 Data acquisition

Transcranial ultrafast ultrasound image acquisition was performed using the Iconeus One (Iconeus, Paris, France) scanner. Images were acquired using a 15 MHz linear array probe (128 elements, 0.1 mm pitch) placed on intact skull and skin after hair removal and the application of echographic gel. Throughout the experiment, the probe was fixed steadily on a motorized system (Fig. 1). Prior to recordings, the target plane was determined by performing a 3D whole brain fUSI image acquisition for each mouse, and co-registering with a standard Allen Mouse Common Coordinates Framework brain atlas utilizing dedicated software available with the Iconeus system [17]. pD images were acquired with a spatial resolution of 100 µm × 100 µm, 400 µm slice thickness, and 12.8 mm (width) × 10 mm (depth) field of view (FOV). Each image was obtained from 200 compounded frames acquired with a 500 Hz frame rate, with each frame built using 11 tilted plane waves (from -10° to +10°, increment by 2°) acquired at 5500 Hz pulse repetition frequency. fUSI recording sessions were performed using real-time continuous acquisition, with successive blocks of 400 ms, separated by 600 ms pause, generating a frame rate of 1 Hz. In the current study, we used 2 min of fUSI recordings prior to MK-801 drug or saline injection for evaluating the image registration techniques.

**Figure 1.**
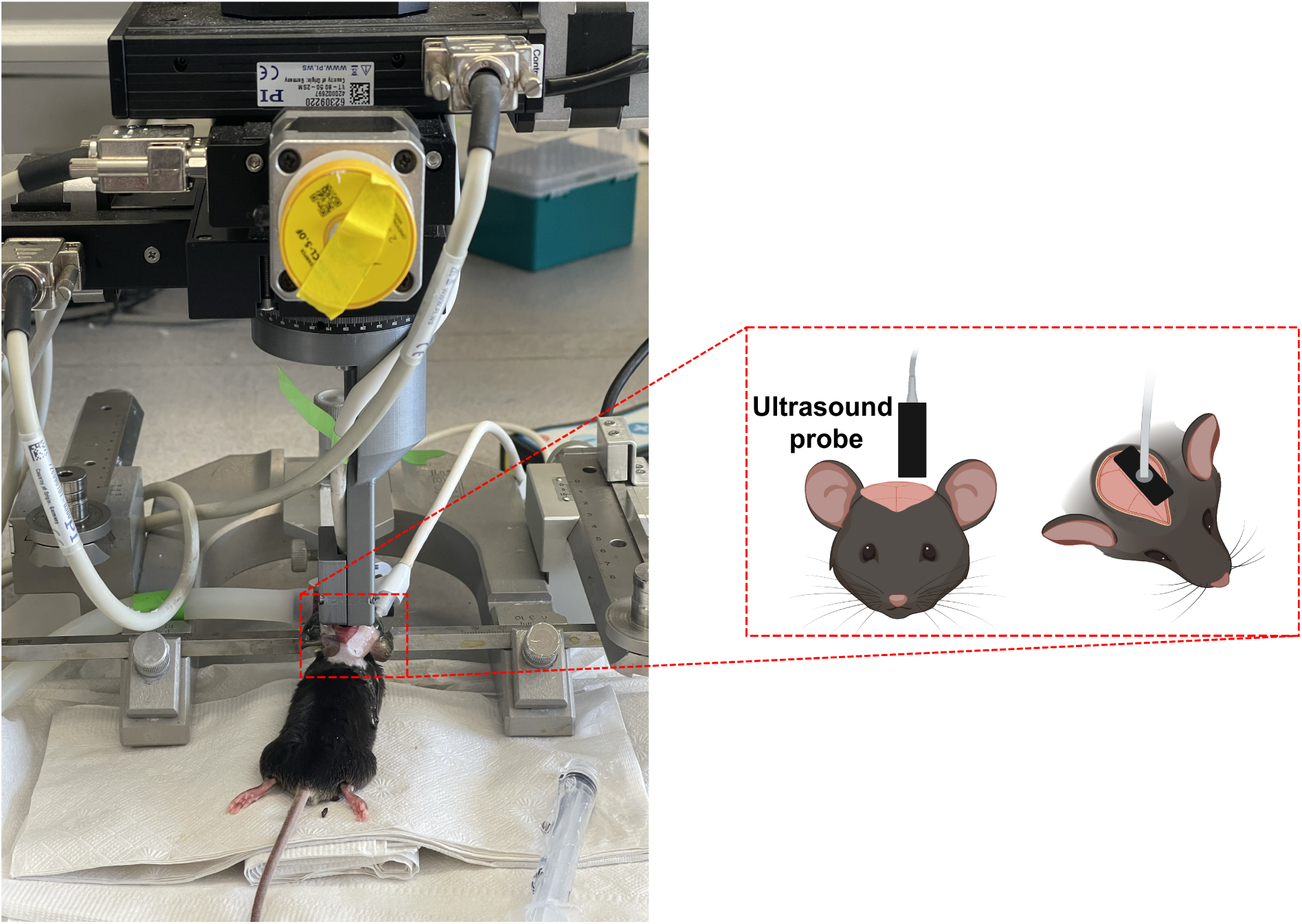
Experimental setup and recordings. An illustrative diagram showing the motorized acquisition system used for 2D fUSI scanning in a mouse brain.

### 2.3 Data pre-processing

We corrected for high frequency fluctuations in the pD signal by applying a low-pass filter with normalized passband frequency of 0.02 Hz, with a stopband attenuation of 60 dB that counterbalances the delay caused by the filter, in order to remove noise artifacts.

### 2.4 Registration preparation

We selected 14 animals in which their average pD vascular maps show the most details compared to the other animals (see Fig. S1 in the supplementary materials). These were designated as reference mice. For each reference mouse, we computed the mean of the first ten images to use as the fixed image (reference image). For each round of registration, one of these 14 fixed images were chosen, and the recordings from the remaining 81 mice were registered to the fixed image using each of the registration techniques.

### 2.5 registration techniques

We employed 6 image registration techniques, using 14 distinct reference mice, and applied these methods to a total of 82 mice. An overview of the methods is shown in Table 1.

**Table 1.** registration techniques utilized in this study.

| Name | Platform | Transform | Metric | Optimization Method | Number of Parameters |
| --- | --- | --- | --- | --- | --- |
| NoRMCorre | Matlab | Rigid/piecewise rigid (non-rigid) | Normalized cross-correlation | One-pass (optimization methods could be added) | 7(rigid)/14(non-rigid) |
| Elastix | Matlab, python | Rigid (non-rigid available) | Mutual information (other metrics available) | Adaptive stochastic gradient descent (others available) | 25 |
| Imregtform | Matlab (built-in) | Rigid/Affine/Similarity | Mutual information (mean squares available) | (1+1) evolutionary optimization strategy (gradient descent available) | 9 |
| Imregdemons | Matlab (built-in) | Non-rigid | Sum of squared difference | Gradient descent | 3 |
| Imregdeform | Matlab (built-in) | Non-rigid | Mutual information + a regularization term | Gradient based optimization | 4 |
| Optic flow | Matlab, python | Non-rigid | Sum of squared difference (ILK)/Total variation and L1 norm (TV-L1) | Gradient based optimization (ILK)/a primal-dual algorithm (TV-L1) | 4(ILK)/6(TV-L1) |

#### 2.5.1 NoRMCorre

NoRMCorre has been commonly used in solving large scale image registration problems in functional neuroimaging [18]. It operates by splitting the field of view (FOV) into overlapping spatial patches (i.e., frames) along all directions. The patches are registered at a sub-pixel resolution for rigid translation against a regularly updated template. To enable a fair comparison between different registration techniques, we deactivated the template updating function, ensuring that the same reference image was consistently used as the template for each round of registration. The estimated alignments are subsequently up-sampled to create a smooth motion field for each image that can efficiently approximate non-rigid artifacts in a piecewise-rigid manner. Note that the NoRMCorre technique could operate in both rigid and non-rigid (piecewise-rigid) fashion, both of which was deployed in our study.

#### 2.5.2 Elastix

*Elastix* toolbox is a modular software package widely used for intensity-based image registration of medical images [19]. It applies a transformation matrix to pixels of moving images (pre-registration image), mapping them onto a fixed image, forming a coordinate map for each pixel. In practice, *Elastix* loops over all pixels in the fixed image, computes the mapped position, interpolates the moving image at the new location, and fills these values into the resulting registered image.

*Elastix* utilizes various key components, such as transformation model, sampler, metric, optimizer, interpolator, and image masks. All these components are adjustable. In this study, we opted to use the Euler transformation (i.e., a rigid transformation) as our transformation model, in which case the image is considered a rigid body subject to translations and rotations. To measure the transformation efficiency, we initially utilized a sampler strategy to select locations (i.e., pixels) in the fixed image for the metric to evaluate. The most straightforward strategy is to use all pixels from the fixed image, with the obvious downside that it is time-consuming. On the other hand, the random sampler strategy randomly selects a subset of pixels in the fixed image. Every pixel has an equal probability to be selected and a sample may be selected more than once. In our study, we used an extension of the random sampler strategy, named “Random Coordinate” that is not limited to pixel positions, but also samples positions between pixels.

The next step is to select a similarity metric that measures the degree of similarity between the fixed and the moving images. Among different metrics, we selected the Normalized Mutual Information (NMI) metric component that measures the ratio of the images’ marginal histogram (pixel intensity distribution) and their joint histogram:

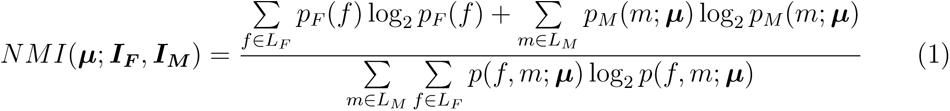

Where ***µ*** is the transformation parameter, ***I_F_*** and ***I_M_*** are the fixed and moving images, respectively, *p* is the joint probability, and *p_M_* and *p_F_*are the marginal probabilities calculated by summing *p* over *m* and *f* . The measure produced by the NMI metric provides a criterion that is optimized using the Adaptive Stochastic Gradient Descent (ASGD) method, which has good convergence properties and is relatively fast in combination with the random sampling strategy. ASGD estimates the optimal transform parameters by enabling the transformation matrix to be adjusted in each iteration. This is accomplished by taking adaptive step sizes towards the direction that minimizes the NMI function (i.e., cost function *C*) as shown in Eq. 2.

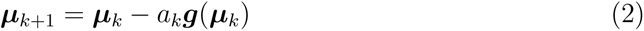

where 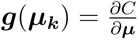, which represents the gradient of the cost function *C* with respect to the transformation parameter ***µ*** evaluated at the current position ***µ_k_***, and *a_k_* is the gain factor. Moreover, a transformed point is generally mapped to a non-grid position, which is not precisely aligned with a pixel’s exact grid position. During optimization, the transformed point requires interpolation to evaluate the image intensity. A linear B-spline interpolation was performed using a 1st-order B-spline function. This function calculates a weighted average based on the surrounding pixels. The weight assigned to each surrounding pixel is determined by its distance to the point where the interpolated value is needed. Note that we opted for 1st-order B-spline interpolation because it offers a good trade-off between quality and computation time, as well as to prevent the overshoot characteristic often associated with higher-order B-spline interpolations.

Finally, a binary image mask was implemented to specify a region of interest used for registration. Using the gray threshold values, the corresponding binary images were constructed for both moving and fixed images, indicating the registered region. Regions with “0” value were designated as “false”, and regions with “1” value were “true” or pixels of interest.

#### 2.5.3 Imregtform

We utilized *Imregtform*, a function native to the Image Processing Toolbox in Matlab that performs intensity-based image registration. This technique operates on the principle of identifying an optimal geometric transformation to align the moving image to a fixed image [20]. To achieve this, we implemented a (1+1) evolutionary optimization strategy that draws inspiration from the basic principles of natural selection and mutation [21]. This approach is a simple yet effective form of evolutionary algorithm used for optimization problems. In this method, one single parent produces one single offspring in each generation through mutation. If the offspring’s fitness is better or equal to the parent’s, it replaces the parent in the next generation [22]. We applied this strategy using a *OnePlusOneEvolutionary* object to register moving images to fixed images.

The *Imregtform* function allows the use of various transformation types to adjust the spatial relations of the images under consideration. These include translation (shift in X and Y coordinates), rigid (translation plus rotation), affine (includes scaling and shearing), and similarity transformations (preserves shape, includes rotation, translation, and scaling) [20]. In this study, we specifically implemented rigid, affine and similarity transformations for the registration process. Rigid transformation preserves the distances between every pair of points in the moving image. It is especially beneficial in scenarios where rotation and translation are the only changes between the fixed and moving images [23]. Affine transformation allows shearing and scaling besides rotation and translation, offering more degrees of freedom to handle more complex deformations [23]. Lastly, similarity transformation, a subcategory of the affine transformation that specifically preserves the shape of objects, is advantageous when registering images of similar objects taken from varying perspectives or scales.

#### 2.5.4 Imregdemons

We also utilized *Imregdemons*, a function that is available in the Image Processing Toolbox in Matlab, to implement a variation of the “Demons” algorithm proposed by J.P.Thirion [24, 25]. This technique adopts the concept of diffusing models to perform image-to-image matching. The major principle of the Demons algorithm lies in its approach to estimate a displacement field that maps the moving image onto the fixed image. It works under the assumption that the intensity of a given point should be the same in both the moving and fixed images, and if it is not, a “force” or “demons” is created that pushes the moving image towards the fixed image. This idea is similar to how Maxwell’s approach resolved the Gibbs paradox in thermodynamics. The exerted forces were conceptualized from the principles of optical flow equations [26], and is proportional to the intensity difference and the gradient of the fixed image.

The displacement resulting from the application of the force by the demons is given by the equation:

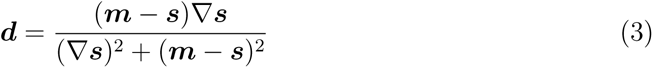

where ***d*** is the displacement during one iteration step, ***m*** and ***s*** are the intensities of the moving and fixed images, respectively, and ∇***s*** is the gradient in each nodal point of the fixed image [25]. The algorithm operates by alternately calculating these forces and performing regularization through elementary Gaussian smoothing. The Demons algorithm was then further refined to enforce diffeomorphic transformations [27], which is also implemented by the *Imregdemons* function.

In this study, the default settings for Matlab *Imregdemons* were used, which includes 3 pyramid levels, 100 iterations, and a Gaussian smoothing with a standard deviation of 1 at each iteration to regularize the accumulated field (see registration techniques parameters in the supplementary materials for more details).

#### 2.5.5 Imregdeform

*Imregdeform* is part of the Matlab Image Processing Toolbox (since 2022b) [28]. It utilizes a parametric approach for image registration with total variation regularization. This approach is characterized by the incorporation of an isotropic total variation [29] and an efficient local correlation coefficient (LCC), which is defined as the combined weighted correlation coefficients of image intensity levels, computed over n-dimensional patches centered at each pixel [30]. The isotropic total variation is defined by the following equation [30], :

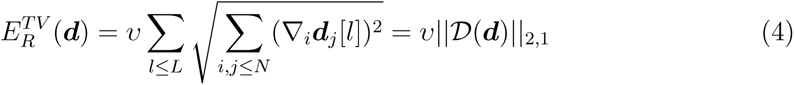

where ***d*** is the displacement field, ∇*_i_****d****_j_*[*l*] denotes the spatial gradient of the *j*-th displacement component at location *l*, and *υ* is the pixel volume. The term *||D*(***d***)*||*_2,1_ is the 2, 1-norm of the displacement gradient matrix *D*(***d***). The displacement gradient matrix *D*(*d*) is defined as [30]:

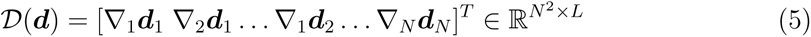

with each term representing the spatial gradient for each component of the displacement field in each dimension.

This method provides an efficient solution scheme via the alternating directions method of multipliers (ADMM), enabling better convergence in practice than conventional comparable methods. In this study, the default settings for *Imregdeform* were applied, which includes 3 pyramid levels, a grid spacing value of [4 4], a pixel size of [1 1], and a weighting factor of 0.11 for grid displacement regularization (see registration techniques parameters in the supplementary materials for more details).

#### 2.5.6 Optical Flow

The optical flow algorithm is a method for estimating the displacement of an image region from one image to the next. We opted to try two variants of the method; the original TV-L1 [31], and the computationally cheaper Lukas Kanade (ILK) method, which is an approximation method that prioritizes speed over accuracy [32]. Although optical flow is typically used for tracking the motion of objects between consecutive frames of a video, in the context of this study, we used only a single fixed image for an entire sequence of moving images.

The algorithm works by analyzing the changes in pixel brightness between two images and estimating the displacement. The displacement is computed based on known intensity gradients of the image in the specific region, and can be described by a vector (**u**, **v**), where **u** and **v** denote the displacement in the x and y directions, respectively. The change in brightness in the x and y directions are denoted as *I_x_***u** and *I_y_***v**, respectively, and the change of brightness over time is given by *I_t_*. These changes can be modeled by the constraint equation Eq. 6 that determines the change in the intensity of a pixel from one image to its corresponding position in the next image.

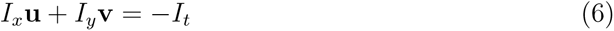

which can be rewritten as:

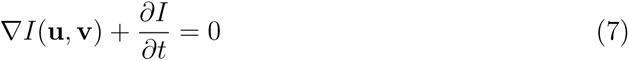

The constraint equation relies on two assumptions: a) the objects in the image do not change their brightness over time, and b) they are moving at a reasonably slow speed between images.

Fundamentally, the two optical flow variants utilize the same constraint equation, but they diverge in their respective methods of solving it. The original TV-L1 variant addresses the constraint equation by selecting the (**u**, **v**) that minimizes the error, as defined by the Horn-Schunck equation:

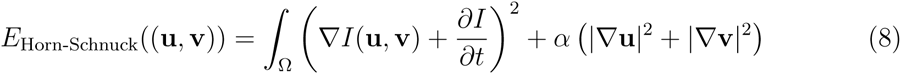

The ILK method proposes an alternative solution to the constraint equation. Instead of treating each pixel individually, the ILK method formulates a system of equations from a kernel of pixel**s**, and then derives the least squares solution to average the estimation across the entire kernel. Consider a kernel of size *n*, centered around pixel (**x**, **y**). The constraint equation can be applied to each pixel within this kernel, allowing every pixel to be represented by its corresponding equation:

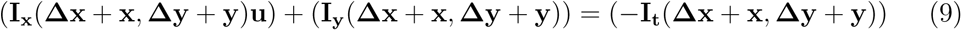

Accounting for all pixels within the kernel, we can represent the left side of the resulting system of linear equations as the *n ×* 2 matrix **S**, and the right side as the vector **t**, resulting in the matrix equation:

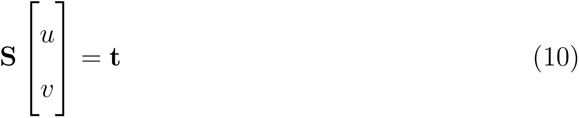

With this equation, the displacement vector (**u**, **v**) can be estimated as:

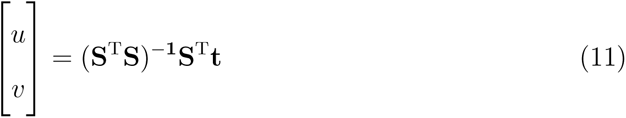

Typically, utilizing a smaller kernel allows for finer image manipulation, which often yields a higher degree of similarity between the registered and fixed images. We chose the default value, which is a uniform kernel with a radius of 7 performed over a single iteration.

The two optical flow variants represent distinct methods of creating a vector field. This field is the resultant matrix of vectors (**u**, **v**), determined for each individual pixel. The final transformed image is produced by shifting every pixel as instructed by the associated vector.

### 2.6 Evaluation measures

We used a variety of metrics to evaluate the performance of the image registration techniques.

#### 2.6.1 Dice Similarity Coefficient

We computed the dice similarity coefficient (DSC) for comparing the similarity between the moving and registered images with the reference pD vascular map (fixed image). For a pair of 2-dimensional images, the DSC is defined as follows:

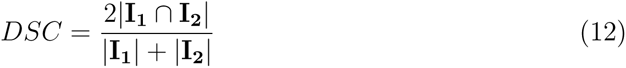

where **I_1_** and **I_2_** represent two binary images, ‘*∩*’ represents the intersection operation, and ‘| |’ represents the size of a set. In other words, DSC is defined as twice the area of overlap between two images divided by the total number of pixels in both images. Before computing the DSC, we use the imbinarize function in Matlab to binarize the 2-dimensional surfaces by thresholding using Otsu’s method [33]. Briefly, it chooses a global threshold that minimizes the inter-class variance of the thresholded black and white pixels. Then we compute the DSC between each moving and registered images with the fixed image.

#### 2.6.2 Mutual information

Mutual information (MI) quantifies the statistical dependence or information shared between the intensity values of pixels in two images [34]. It is defined as the sum over all intensity values of the joint probability of the intensities in the two images multiplied by the logarithm of the ratio of the joint probability to the product of the marginal probabilities:

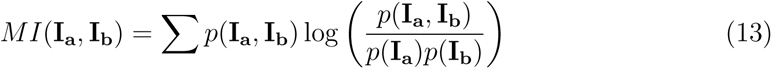

Where **I_a_** and **I_b_** represent the pixel intensities of two images, *p*(**I_a_**, **I_b_**) is the joint probability of the intensities in images **I_a_** and **I_b_**, and *p*(**I_a_**) and *p*(**I_b_**) are the marginal probabilities of the intensities in images **I_a_** and **I_b_**, respectively. We computed the MI between each moving and registered images with the fixed image.

#### 2.6.3 Normalized cross-correlation

Normalized cross-correlation is robust to changes in brightness and contrast in the images being compared, which is used to determine the degree of alignment between two different images. It is defined as the sum over all pixels of the product of the intensities in the two images, normalized by the product of the standard deviations of the intensities:

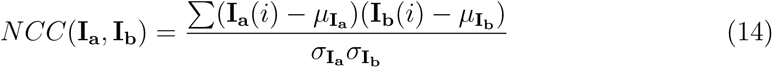

Where **I_a_**and **I_b_** are the pixel intensities of two images, *i* is the index for the pixels in the images, *µ***_I_** and *µ***_I_** are the mean signal intensities in **I_a_**and **I_b_**, and *σ***_I_** and *σ***_I_** are the standard deviations of the intensities in **I_a_** and **I_b_**, respectively.

We computed the normalized cross-correlation using two different methods: First, we used the *normxcorr*2 function built in Matlab to directly compute the 2D cross-correlation between the mean of each moving and registered image and each fixed image, and measured the peak value, peak location and width. Second, we computed the normalized cross-correlation between each moving and registered image and each fixed image assuming that the images are already aligned without considering any shifts or lags.

#### 2.6.4 Mean Squared Error

Mean squared error (MSE) quantifies the average squared difference between the corresponding pixels between images. The MSE is defined as:

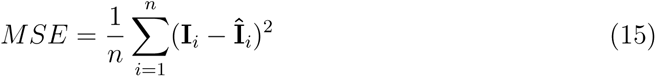

Where *n* is the total number of pixels, **I** represents the pixel intensities of one image and **^^^I** represents the corresponding pixel intensities of another image. We utilized the *immse* function in Matlab to compute the MSE between each moving and registered image and the fixed image.

## 3 Results

### 3.1 General assessment of fUSI image registration

We evaluated 6 distinct image registration techniques (refer to Table 1 for details). For NoRMCorre, both rigid and piecewise-rigid transformations were performed. For *Imregtform*, affine, rigid, and similarity transformations were separately performed. We evaluated these techniques using fUSI recordings from 82 anesthetized mice. We selected 14 animals with high quality vascular maps to serve as our reference group. We then computed the mean vascular map from the first 10 images from each of these mice to generate the fixed image for each animal (see Fig. S1 in the supplementary materials). Fig. 2 depicts the general performance of image registration techniques using two example mice. One is selected as the reference animal and the other is chosen for registration. The goal is to perform ultrafast Doppler vascular-to-vascular image registration between them. The top green-magenta panel of the figure represents the initial misalignment between the average vascular maps of the two animals before registration. The fixed image from the reference animal is displayed in green, while the other animal is considered to be the moving image (magenta) that we want to register to the fixed image using the set of image registration techniques described before. The bottom panels of Fig. 2 illustrate the overlaid vascular maps of the two animals, with the moving image registered to the fixed image using the image registration techniques utilized in the current study.

**Figure 2.**
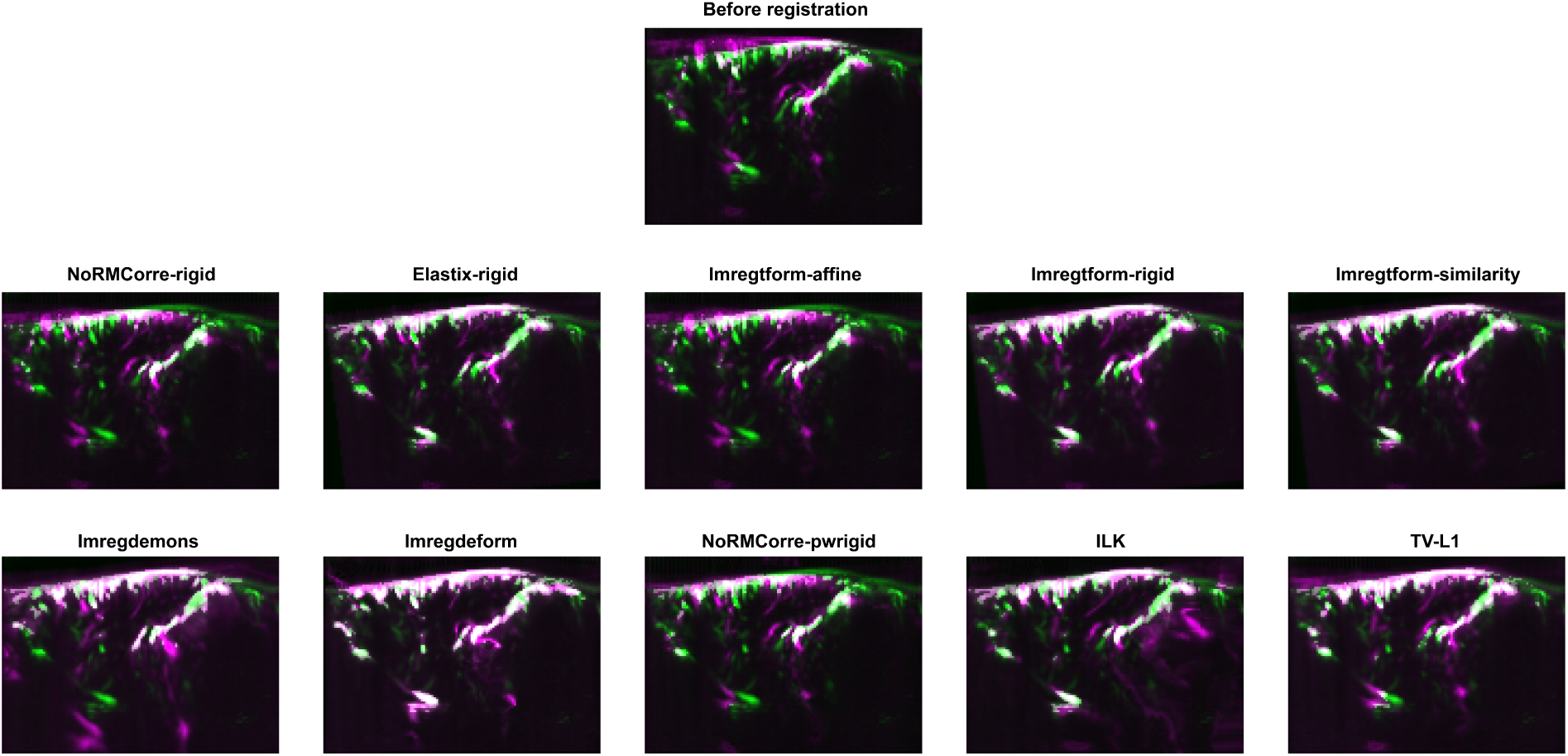
Visual assessment of fUSI image registration. Typical example of fUSI images obtained in a pair of acquisitions from two different mice acquired at different time points, before registration (top) and after registration (bottom). Green: fixed image. Magenta: moving/registered image.

### 3.2 Quantitative evaluation of registration techniques

We quantified the performance of the image registration techniques by computing the 2D normalized cross-correlation (NCC) between the average fixed image and the average moving and registered images in the two space directions (Fig. 3). The NCC between the fixed and the registered images shows a stronger (value range from 0.45 to 0.66) and sharper (width at half-maximum range from 5 to 7 pixels for x direction, and 3 to 6 pixels for y direction) peak located in the center, compared to the NCC between the fixed and the moving images both in x and y directions (value = 0.28 and width at half-maximum = 12 pixels for x direction and 9 pixels for y direction). As reference, autocorrelation (same fixed image) yields a width at half-maximum = 5 pixels for x direction and 3 pixels for y direction. Among the different registration techniques, *Imregdeform* exhibits the highest peak in comparison to all other non-rigid and rigid methods, closely followed by the non-rigid method *imregdemons*. The *imregdemons*, NoRMCorre’s piecewise-rigid method, *Imregtform* (for similarity, rigid, and affine transformation), the *Elastix*-based rigid method, and *TV-L1* showed similar NCC peaks and widths, while the *ILK* and the rigid method for NoRMCorre exhibited the lowest NCC values. Nevertheless, for these two techniques, the NCC peak was still higher and sharper than that of the moving images (black discontinuous trace).

**Figure 3.**
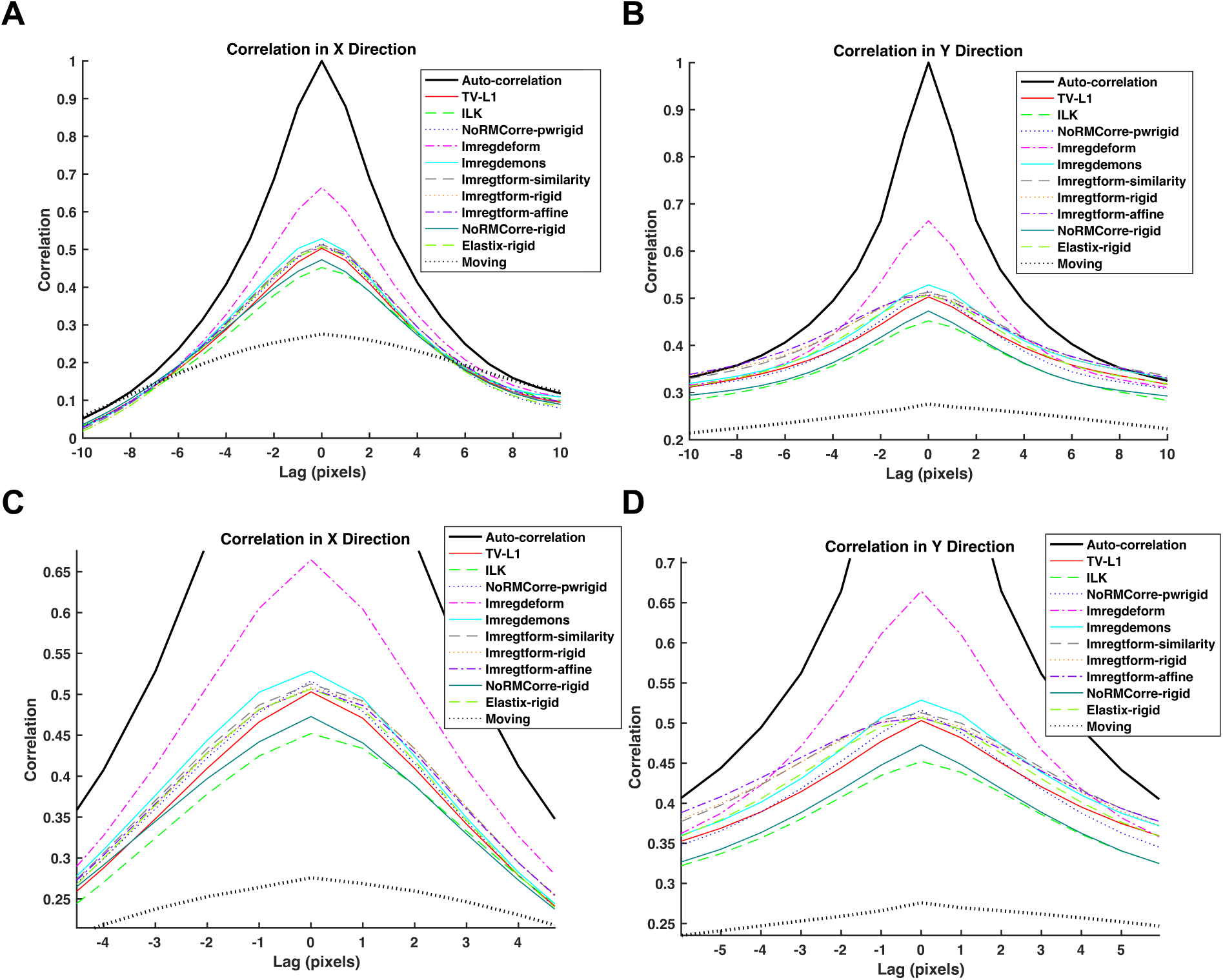
Normalized cross-correlation analysis between fixed image and moving/registered images. (A) Normalized cross-correlation plot between the fixed image and moving/registered images in the x direction. Fixed image auto-correlation is also shown. (B) Same as (A) but for the y direction. (C) and (D) are zoomed in versions of (A) and (B), respectively.

To further evaluate the performance of the image registration techniques, we computed the normalized cross-correlation values for each slice within the moving and registered images relative to the fixed image, without considering any shifts or lags (as shown in Fig.4A). The results were consistent with our previous findings showing that the cross-correlation increased after registration, with *Imregdeform* significantly outperforming all other rigid and non-rigid registration techniques (One-way ANOVA, p<0.001). We also computed three additional metrics to better evaluate the performance of the image registration techniques. In particular, we computed the DSC, which is a widely accepted metric for assessing the similarity or overlap between two sets (see Materials and Methods section for more details), the results are shown in Fig. 4B. The DSC ranges from 0 (no overlap) to 1 (perfect overlap). Mirroring the NCC results, *Imregdeform* excelled (one-way ANOVA, p<0.001) by generating registered images that exhibits the highest DSC in comparison to the fixed image, followed by *Imregdemons* and *Imregtform* (affine and similarity transformation), with no significant difference between these two techniques.

**Figure 4.**
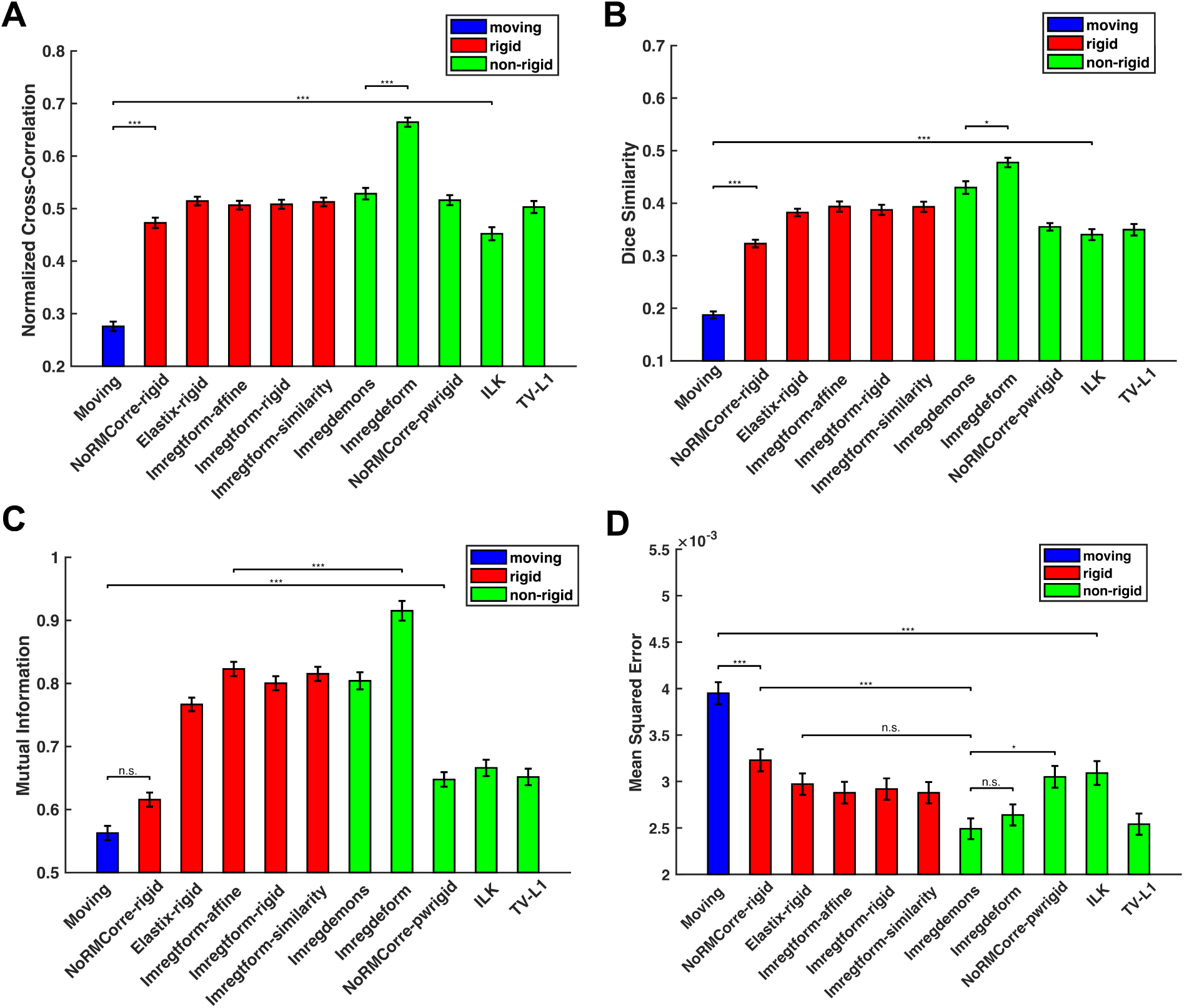
Quantitative evaluation of registration techniques. (A) Quantification of normalized cross-correlation across 14 fixed images. Color code: Blue: Pre-registration (moving) images. Red: Registered images for rigid techniques. Green: Registered images for non-rigid techniques. (B)-(D): Quantification of Dice similarity, mutual information and mean squared error across 14 fixed images for moving and registered images, respectively.

We also computed the MI between each image and the fixed image (Fig. 4C). MI is an entropy-based metric that measures the dependency between the moving/registered and fixed images. Higher MI values indicate better registration performance. For MI, *Imregdeform* remains the best technique (one-way ANOVA, p<0.001), followed by *Imregtform* and *Imregdemons* (no significant difference between these two). Notably, there is no significant difference between the moving and registered images with respect to MI when using the NoRMCorre (rigid) technique (one-way ANOVA, p=0.081) Finally, we computed the mean squared error (MSE) between each moving/registered image and the fixed image (Fig. 4D). The MSE quantifies the average squared difference between pixel intensities of the moving/registered and fixed images (see Methods section for more details). Lower MSE values indicate better registration performance. The results show that MSE significantly decreases following all rigid and non-rigid image registration techniques. Interestingly, *Imregdemons* exhibits the lowest MSE (although only significantly lower than NoRMCorre and *ILK*), The methods *Imregdeform*, *TV-L1*, *Imregtform*, and *Elastix* showed similar performance to *Imregdemons*, with no significant differences observed among them. This seems to be reasonable since *Imregdemons* adopts the sum of squared differences as the metric for registration.

## 4 Discussion

Recent advances in neuroimaging technology have significantly contributed to a better understanding of human brain organization, as well as the development and application of more efficient clinical programs. A key element in neuoroimaging is the process of image registration, a technique that facilitates the alignment of multiple brain images taken at different time points and/or from different subjects [35]. This alignment enables more robust comparisons, correlation and integration of information across these images, contributing significantly to diagnosing, monitoring, and treatment planning procedures in numerous medical contexts [36].

FUSI is a novel avenue in neuroimaging that holds substantial promise for future pre-clinical and clinical applications. Unlike traditional ultrasound imaging, fUSI is based on ultrafast Doppler sequences, which is capable of visualizing not only the vascular anatomy of the brain, but also hemodynamic changes of 1 mm/s or higher as seen in arterioles *>* 10 µm, and thus to perform imaging of localized brain activity through neurovascular coupling [2, 3, 10]. In comparison to other imaging modalities like fMRI, fUSI offers distinct advantages including high sensitivity to slow blood flow, real-time imaging acquisition and superior spatiotemporal performance [37]. The fUSI scanner is like any clinical ultrasound machine, making the unit freely mobile between different settings and negative the need for extensive infrastructure inherent to fMRI. All these advantages open up opportunities for point-of-care applications and revolutionizing bedside monitoring in clinical settings.

However, an integral aspect which hasn’t been thoroughly addressed in fUSI is image registration. Therefore, the preponderance of fUSI studies are restricted to either averaging activity within selected ROIs or ROI-ROI functional connectivity analysis. Given the variability inherent in fUSI due to factors such as variability in anatomical structures across animals, operator-dependency and fluctuations in image quality, a standardized registration pipeline is crucial to allow for accurate comparison and analysis of images. Image registration will also enable more consistent tracking of functional changes over time, further enhancing the diagnostic and monitoring capabilities of fUSI.

In the current study we evaluate a number of image registration techniques, which have been widely used in the neuroimaging literature, to align 2-dimensional brain fUSI datasets acquired from different mice while they were under anesthesia. We found that the *Imregdeform* technique outperforms all other methods under almost all metrics, followed by *Imregdemons* and *Imregtform*. The only exception is when MSE metric is used to evaluate the performance of the methods. In this case, *Imregdemons*, *Imregdeform*, *TV-L1*, *Elastix* and *Imregtform* perform similarly well for aligning the 2-dimensional fUSI brain images. These findings suggest that *Imregdeform* is the best non-rigid image registration technique to align 2-dimensional brain fUSI data across different mice. However, if spatial distortions during the registration process is of concern, rigid image registration techniques such as *Imregtform* or *Elastix* could be considered.

Although our study compares for the first time different image registration techniques in fUSI brain datasets, it has a number of limitations. Firstly, the animals used in this study are from the strain C57BL/6 and aged between 8-12 weeks old. It would be interesting to compare and evaluate the image registration techniques using mice of different ages, pathologies, or different animal models (e.g., rats, non-human primates), which could yield more diverse brain and vasculature shapes across subjects.

Furthermore, in the current study we did not optimize the parameters for each image registration technique. Instead, we used the generic parameters proposed by these methods for aligning the fUSI brain datasets, unless the default parameters have to be changed. For instance, we had to change the default parameters in the *Imregtform* technique, otherwise a warning signal would be displayed stating that registration failed because optimization diverged. As such, the post-registration results might not fully reflect the optimal capabilities of each respective technique. On the other hand, optimizing some of these methods is a challenging procedure that is beyond the scope of the current study, which aims to establish an initial set of guidelines for selecting image registration techniques for fUSI brain datasets. We anticipate that our findings can serve as a foundation for further fine-tuning and advancements in this area of research.

Additionally, although the vast majority of fUSI studies acquire 2-dimensional brain imaging datasets, recent studies attempted to tackle the challenging whole-brain 3-dimensional fUSI using either moving linear arrays [38] (similar to the array used in our study), matrix arrays [39, 40] or raw column arrays (RCAs) [41]. The presented image registration techniques can be extended to align 3-dimensional fUSI datasets, although we need to re-evaluate their performance.

Overall, our study presents a comparison among well-established image registration techniques in computer vision to align a dataset of 2-dimensional fUSI brain recordings from 82 mice. The results showed that the non-rigid image registration technique *Imregdeform* outperforms all alternative methods across almost all evaluation metrics. This revelation is of considerable importance, as it can standardize the image registration protocol for 2-dimensional fUSI brain datasets, enhancing the consistency and reliability of results across studies. This progression is of high significance to the neuroscience research community, as it could facilitate more precise investigations using the fUSI technology in pre-clinical studies.

## CRediT author statement

**Shan Zhong**: Methodology, Software, Formal analysis, Visualization, Writing-Original Draft. **Kofi Agyeman**: Methodology, Software, Writing-Original Draft. **Shanze Syed**: Software, Writing-Original Draft. **Richard Tobing**: Software, Writing-Original Draft. **Wooseong Choi**: Data acquisition. **Charles Liu**: Writing-Reviewing and Editing, Funding acquisition. **Darrin Lee**: Writing-Reviewing and Editing, Funding acquisition. **Vassilios Christopoulos**: Conceptualization, Methodology, Supervision, Writing-Reviewing and Editing, Funding acquisition.

## Acknowledgments

Research reported in this publication was supported by the Hellman Foundation, Brain and Behavior Research Foundation and USC Neurorestoration Center. The funders had no role in study design, data collection and analysis, decision to publish, or preparation of the manuscript.

## References

1. M. Tanter and M. Fink. Ultrafast imaging in biomedical ultrasound. *IEEE Transactions on Ultrasonics*, Ferroelectrics, and Frequency Control, 61(1):102–119, 2014.

2. E. Macé, G. Montaldo, I. Cohen, M. Baulac, M. Fink, and M. Tanter. Functional ultrasound imaging of the brain. Nature Methods, 8(8):662–664, 2011.

3. E. Macé, G. Montaldo, B.-F. Osmanski, I. Cohen, M. Fink, and M. Tanter. Functional ultrasound imaging of the brain: theory and basic principles. *IEEE Transactions on Ultrasonics*, Ferroelectrics, and Frequency Control, 60(3):492–506, 2013.

4. J. Bercoff, G. Montaldo, T. Loupas, D. Savery, F. Mézière, M. Fink, and M. Tanter. Ultrafast compound doppler imaging: Providing full blood flow characterization. IEEE transactions on ultrasonics, ferroelectrics, and frequency control, 58(1):134–147, 2011.

5. B.F. Osmanski, C. Martin, G. Montaldo, P. Lanièce, F. Pain, M. Tanter, and H. Gurden. Functional ultrasound imaging reveals different odor-evoked patterns of vascular activity in the main olfactory bulb and the anterior piriform cortex. Neuroimage, 95:176–84, 2014.

6. B.-F. Osmanski, S. Pezet, A. Ricobaraza, Z. Lenkei, and M. Tanter. Functional ultrasound imaging of intrinsic connectivity in the living rat brain with high spatiotemporal resolution. Nature Communications, 5(1):5023, 2014.

7. L.-A. Sieu, A. Bergel, E. Tiran, T. Deffieux, M. Pernot, J.-L. Gennisson, M. Tanter, and I. Cohen. Eeg and functional ultrasound imaging in mobile rats. Nature Methods, 12(9):831–834, 2015.

8. A. Dizeux, M. Gesnik, H. Ahnine, K. Blaize, F. Arcizet, S. Picaud, J.A. Sahel, T. Deffieux, P. Pouget, and M. Tanter. Functional ultrasound imaging of the brain reveals propagation of task-related brain activity in behaving primates. Nature Communications, 10(1):1400, 2019.

9. S.L. Norman, D. Maresca, V.N. Christopoulos, W.S. Griggs, C. Demene, M. Tanter, M.G. Shapiro, and R.A. Andersen. Single-trial decoding of movement intentions using functional ultrasound neuroimaging. Neuron, 109(9):1554–1566, 2021.

10. M. Imbault, D. Chauvet, J.-L. Gennisson, L. Capelle, and M. Tanter. Intraoperative functional ultrasound imaging of human brain activity. Scientific Reports, 7(1):7304, 2017.

11. C. Demene, J. Baranger, M. Bernal, C. Delanoe, S. Auvin, V. Biran, M. Alison, J. Mairesse, E. Harribaud, M. Pernot, and M. Tanter. Functional ultrasound imaging of brain activity in human newborns. Science translational medicine, 9(411):eaah6756, 2017.

12. B. Vidal, M. Droguerre, M. Valdebenito, L. Zimmer, M. Hamon, F. Mouthon, and M. Charvériat. Pharmaco-fus for characterizing drugs for alzheimer’s disease - the case of thn201, a drug combination of donepezil plus mefloquine. Front Neurosci., 14:835, 2020.

13. C. Rabut, J. Ferrier, A. Bertolo, B. Osmanski, X. Mousset, S. Pezet, T. Deffieux, Z. Lenkei, and M. Tanter. Pharmaco-fus: Quantification of pharmacologically-induced dynamic changes in brain perfusion and connectivity by functional ultrasound imaging in awake mice. Neuroimage, 222:117231, 2020.

14. J.M. Martinez de Paz and E. Macé. Functional ultrasound imaging: A useful tool for functional connectomics? Neuroimage, 245:118722, 2021.

15. T. Deffieux, C. Demené, and M. Tanter. Functional ultrasound imaging: A new imaging modality for neuroscience. Neuroscience, 474:110–121, 2021.

16. L.M. Crown, K. Agyeman, W. Choi, N. Zepeda, S. Siegel, C. Liu, V. Christopoulos, and D. Lee. Frequency- and circuit- specific effects of septohippocampal deep brain stimulation in mice as measured by functional ultrasound imaging. bioRxiv, page 05.21.541598, 2023.

17. Q. Wang, S.L. Ding, Y. Li, J. Royall, D. Feng, P. Lesnar, N. Graddis, M. Naeemi, B. Facer, A. Ho, and T. Dolbeare. The allen mouse brain common coordinate framework: a 3d reference atlas. Cell, 181(4):936–953, 2020.

18. E.A. Pnevmatikakis and A. Giovannucci. Normcorre: An online algorithm for piecewise rigid motion correction of calcium imaging data. Journal of Neuroscience Methods, 291:83–94, 2017.

19. S. Klein, M. Staring, K. Murphy, M.A. Viergever, and J.P. Pluim. Elastix: A toolbox for intensity-based medical image registration. IEEE Transactions on Medical Imaging, 29(1):196–205, 2009.

20. MathWorks. imregtform documentation – matlab, 2023. https://www.mathworks.com/help/images/ref/imregtform.html, Accessed: 2023-07-24.

21. D.E. Goldberg. Genetic Algorithms in Search, Optimization, and Machine Learning. Addison-Wesley, 1989.

22. H-G Beyer and H-P Schwefel. Evolution strategies–a comprehensive introduction. Natural computing, 1:3–52, 2002.

23. D.L.G. Hill, P.G. Batchelor, M. Holden, and D.J. Hawkes. Medical image registration. Physics in Medicine & Biology, 46(3):R1, 2001.

24. J-P. Thirion. Non-rigid matching using demons. In Proceedings CVPR IEEE Computer Society Conference on Computer Vision and Pattern Recognition, pages 245–251. IEEE, 1996.

25. J-P. Thirion. Image matching as a diffusion process: An analogy with maxwell’s demons. Medical Image Analysis, 2(3):243–260, 1998.

26. J.L. Barron, D.J. Fleet, and S.S. Beauchemin. Performance of optical flow techniques. International Journal of Computer Vision, 12:43–77, 1994.

27. T. Vercauteren, X. Pennec, A. Perchant, and N. Ayache. Diffeomorphic demons: Efficient non-parametric image registration. NeuroImage, 45(1):S61–S72, 2009.

28. MathWorks. imregtform documentation – matlab, 2023. https://www.mathworks.com/help/medical-imaging/ref/imregdeform.html, Accessed: 2023-07-24.

29. O. Christiansen, T.M. Lee, J. Lie, U. Sinha, and T.F. Chan. Total variation regularization of matrix-valued images. International Journal of Biomedical Imaging, 2007, 2007.

30. V. Vishnevskiy, T. Gass, G. Szekely, C. Tanner, and O. Goksel. Isotropic total variation regularization of displacements in parametric image registration. IEEE Transactions on Medical Imaging, 36(2):385–395, 2016.

31. J. Sánchez Pérez, E. Meinhardt-Llopis, and G. Facciolo. Tv-l1 optical flow estimation. Image Processing On Line, 3:137–150, 2013. 10.5201/ipol.2013.26.

32. B.D. Lucas and T. Kanade. An iterative image registration technique with an application to stereo vision. In IJCAI’81: 7th International Joint Conference on Artificial Intelligence, volume 2, pages 674–679, 1981.

33. N. Otsu. A threshold selection method from gray-level histograms. *IEEE Transactions on Systems*, Man, and Cybernetics, 9(1):62–66, 1979.

34. F. Maes, A. Collignon, D. Vandermeulen, G. Marchal, and P. Suetens. Multimodality image registration by maximization of mutual information. IEEE Transactions on Medical Imaging, 16(2):187–198, 1997.

35. J.A. Maintz and M.A. Viergever. A survey of medical image registration. Medical Image Analysis, 2(1):1–36, 1998.

36. B. Zitova and J. Flusser. Image registration methods: a survey. Image and Vision Computing, 21(11):977–1000, 2003.

37. T. Deffieux, C. Demene, M. Pernot, and M. Tanter. Functional ultrasound neuroimaging: a review of the preclinical and clinical state of the art. Current Opinion in Neurobiology, 50:128–135, 2018.

38. A. Bertolo, M. Nouhoum, S. Cazzanelli, J. Ferrier, J.C. Mariani, A. Kliewer, B. Belliard, B.F. Osmanski, T. Deffieux, S. Pezet, and Z. Lenkei. Whole-brain 3d activation and functional connectivity mapping in mice using transcranial functional ultrasound imaging. Journal of Visualized Experiments, 168, 2021.

39. C. Rabut, M. Correia, V. Finel, S. Pezet, M. Pernot, T. Deffieux, and M. Tanter. 4d functional ultrasound imaging of whole-brain activity in rodents. Nature Methods, 10:994–997, 2019.

40. C. Brunner, M. Grillet, A. Sans-Dublanc, K. Farrow, T. Lambert, E. Mace, G. Montaldo, and A. Urban. Platform for brain-wide volumetric functional ultrasound imaging and analysis of circuit dynamics in awake mice. Neuron, 108:861–875, 2020.

41. J. Sauvage, J. Porée, C. Rabut, G. Férin, M. Flesch, B. Rosinski, A. Nguyen-Dinh, M. Tanter, M. Pernot, and T. Deffieux. 4d functional imaging of the rat brain using a large aperture row-column array. IEEE Transactions on Medical Imaging, 39:1884–1893, 2020.

